# Deep learning-based chest X-ray age serves as a novel biomarker for cardiovascular aging

**DOI:** 10.1101/2021.03.24.436773

**Authors:** Hirotaka Ieki, Kaoru Ito, Mike Saji, Rei Kawakami, Yuji Nagatomo, Satoshi Koyama, Hiroshi Matsunaga, Kazuo Miyazawa, Kouichi Ozaki, Yoshihiro Onouchi, Susumu Katsushika, Ryo Matsuoka, Hiroki Shinohara, Toshihiro Yamaguchi, Satoshi Kodera, Yasutomi Higashikuni, Katsuhito Fujiu, Hiroshi Akazawa, Mitsuaki Isobe, Tsutomu Yoshikawa, Issei Komuro

## Abstract

Chest X-ray (CXR) is one of the most commonly performed medical imaging tests. Although aging, sex and disease status have been known to cause changes in CXR findings, the extent of these effects has not been fully characterized. Here, we present a deep neural network (DNN) model trained using more than 100,000 CXRs to estimate the patient’s age and sex solely from CXRs. Our DNN exhibited high performance in terms of estimating age and sex, with Pearson’s correlation coefficient between the actual and estimated age of above 0.9 and an area under the ROC curve of 0.98 for sex estimation. The difference between the actual and estimated age is large in CXRs with abnormal findings, suggesting that the estimated age (“CXR age”) can be a biomarker for disease status. Furthermore, by applying our DNN to CXRs of consecutive 1,562 hospitalized heart failure patients, we demonstrated that an elevated CXR age is not only associated with aging-related diseases, such as hypertension and atrial fibrillation, but also a worse outcome of heart failure. Given these results, our new concept “CXR age” serves as a novel biomarker for cardiovascular aging and can help clinicians to predict, prevent, and manage cardiovascular diseases.

## Introduction

Aging is a term used to describe a correlated set of declines in functioning with advancing chronological age. Perceived age, or the estimated age of a person, is a robust biomarker for aging. In clinical practice, physicians unconsciously compare perceived and chronological age. ^1^ Previous clinical studies have revealed that patients with older perceived age, that is, those who look older than their chronological age, have advanced carotid atherosclerosis ^2^, reduced bone mineral density ^3^ and increased mortality ^4^. However, in these studies, perceived age was estimated from facial image of a patient by more than 10 medical professionals and averaged ^4 2 3 5^, so it is not an objective index that can be used in actual clinical practice. In recent years, machine learning-based methods have been developed to estimate the presence of Alzheimer’s disease ^6^ and coronary artery disease ^7^ from facial images of patients. Although perceived age is a useful biomarker for age-related disease and aging, due to privacy and ethical issues, it is difficult to obtain facial images of patients in daily clinical practice.

Because performing a chest X-ray (CXR) is fast and easy, it is one of the most commonly used screening tests for a variety of diseases ^8^. Despite its simplicity and ease of use, CXR provides a lot of information and is pivotal for the diagnosis and monitoring of cardiovascular and pulmonary diseases such as heart failure, aortic dissection, pneumonia, lung cancer, tuberculosis, sarcoidosis, and lung fibrosis ^9^. Although radiological findings of CXR are affected by age ^10^ and sex difference ^11^, few previous studies have demonstrated whether age and sex could be predicted from CXR image ^12, 13^. Several studies have been conducted on automatic diagnosis of CXR; however, it is still difficult to argue that the findings on CXR are not bad for the patient’s age, or that they are trivial findings but abnormal for the patient’s age, or that some findings are normal for male sex.

Recently, deep learning has revolutionized the field of machine learning. Deep neural networks (DNNs) are computational models based on artificial neural networks, consisting of multiple layers that progressively extract higher-level features from raw input. DNNs have been shown to exceed human performance in computer vision and natural language processing tasks ^14^. They have also been applied to the medical field in dermatology, radiology, ophthalmology, and cardiovascular medicine, and have achieved human physician-level performance, for instance, in classifying photographs of skin cancer ^15^, pneumonia detection from CXRs ^16^, diagnosing retinal disease ^17^ and arrhythmia classification from electrocardiograms (ECGs) ^18, 19^. Furthermore, some recent studies have described the possibilities of using DNNs to learn patterns that humans have difficulty in recognizing, such as genetic mutation prediction from histopathology images of lung cancer ^20^, paroxysmal atrial fibrillation pattern detection from normal sinus rhythm ECGs ^21^, cancer treatment response prediction from CT images ^22^, age and sex estimation from ECGs ^23^ and brain age estimation from magnetic resonance imaging (MRI) ^24^. Furthermore, age estimated from ECGs was reported to be associated with comorbidities such as hypertension and diabetes, suggesting the potential of age estimation as a health indicator^23^, and “brain age” estimated from brain MRI is also associated with future progression of dementia^24^. These examples suggest the potential of using a DNN to obtain unseen and useful information from commonly used tests.

We hypothesized that estimated age from CXR using deep learning (“CXR age”) could be an indicator for aging status. In this study, we sought to develop and train DNNs to estimate patients’ age and sex solely from frontal-view CXRs without any additional clinical information and evaluated its estimation performance using a robust and unbiased method. Because CXRs are widely used, we assumed that it can provide great clinical significance if the extent of aging can be estimated from CXRs, “CXR age” could be used as a substitute for perceived age. We explored the clinical implications of “CXR age” by analyzing the relationship between CXR age and CXR findings. We applied the developed DNN to the CXRs of heart failure (HF) patients and examined its relationship with the patient’s background, clinical parameters, and HF outcome.

## Results

### Dataset and Model training

An overview of this study is shown in **Fig. 1**. First, we used the NIH chest X-ray dataset to develop a DNN that estimates the patient’s age and sex from CXR ^25^. This dataset is a large publicly available image dataset containing 112,120 png images of frontal-view CXR from 30,805 unique patients. The dataset also includes metadata containing patients’ age and sex information with finding labels. After removing the age outliers, 63,328 (56%) of the 112,104 X-rays were male CXRs. The ages ranged between 1 and 95 years, with a median age of 49 years and an interquartile range of 35– 59 years (**Supplementary Figs. 1a, d**). We randomly assigned this data to the training, validation, and test data (**Supplementary Fig. 2**).

**Fig. 1.**
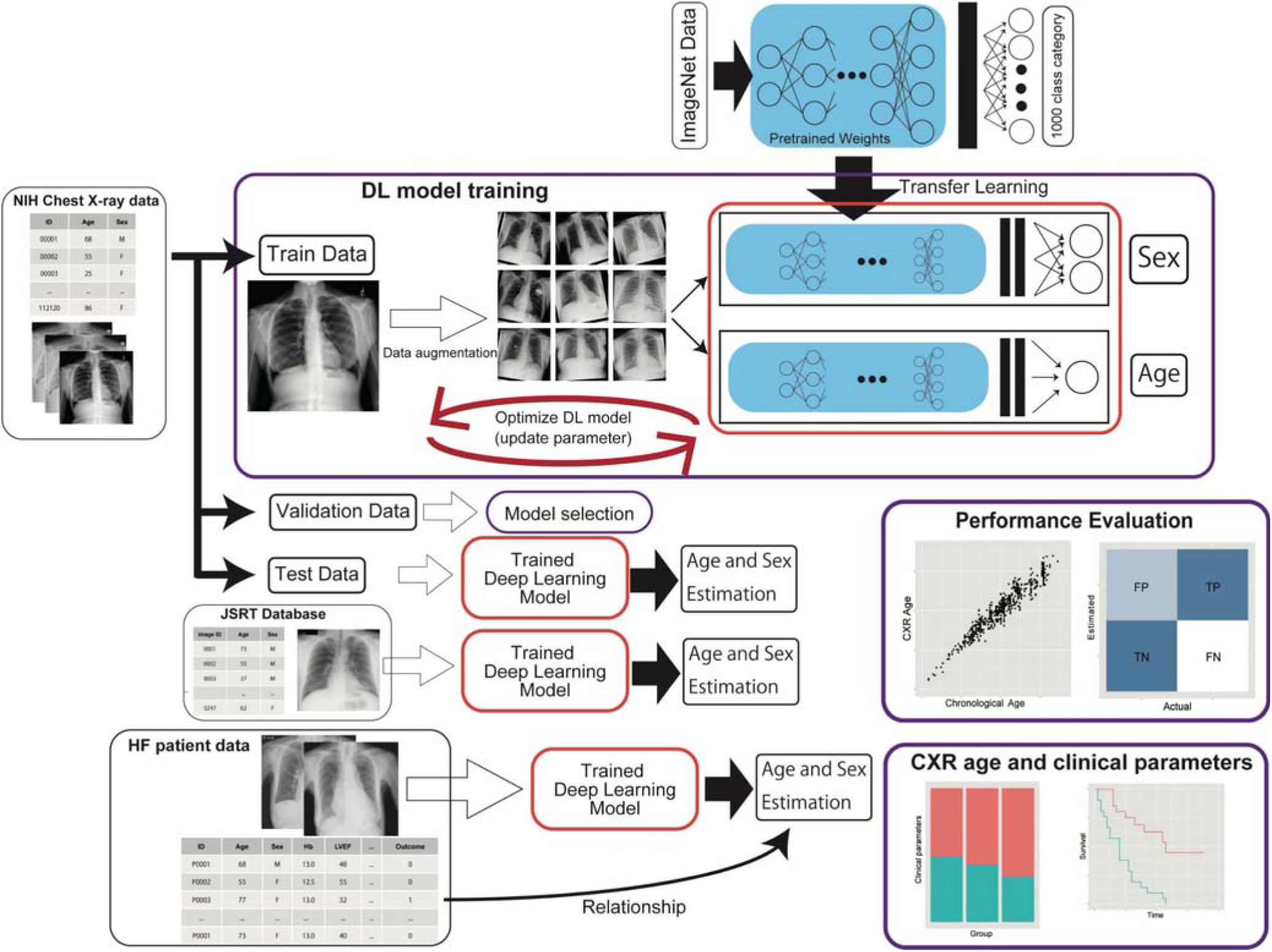
Data usage and overall study framework. The NIH Chest X-ray dataset was randomly divided into training, validation, and test datasets. Our deep neural network (DNN) models were trained to estimate age and sex using the training dataset. The weights of the models were initialized with pre-trained weights on ImageNet data and trained using transfer learning and fine-tuning techniques. Various models with different architectures were separately trained. Validation data was only used to tune the hyperparameters and to select the final model. The accuracy of the deep learning model was estimated using the hold-out test dataset. The independent JSRT dataset was also used to estimate the performance to verify the generalizability of the trained DNN in an independent population. The trained DNN was applied to CXRs of heart failure patients to evaluate the association between the estimated age (CXR age) and various clinical parameters and heart failure.

We applied the transfer learning and fine-tuning techniques to train the DNN. Briefly, these methods utilize a pre-trained DNN to improve the efficiency of the training time and the amount of data used for training. Rather than training the DNN from scratch, the DNN can learn much faster and with significantly fewer training examples by using transfer learning and fine-tuning ^26, 27^. We adopted four commonly used architectures, namely ResNet ^28^, DenseNet ^29^, Inception-v4 ^30^ and SENet ^31^ as pre-trained DNNs. To improve the generalizability of our DNN and avoid overfitting, we applied image augmentation ^32^. After the training, we selected the model with the lowest loss value in the validation dataset as the final model. The metrics of the model with the lowest loss in the validation dataset for each architecture are summarized in **Supplementary Tables 1 and 2**. For age estimation, the SENet-based model yielded the lowest mean squared error (mean squared error: 27.34 (validation dataset)). The Densenet161-based model yielded the lowest binary cross-entropy loss in the validation data (binary cross-entropy: 0.0430 (validation dataset)) in the sex estimation task (**Supplementary Fig. 3**). All the CXR images in the holdout test dataset were used to measure the performance of the model. For age estimation, the estimated age showed a very strong significant correlation with chronological age (Pearson’s r: 0.961, p < 2.2 × 10^-323^) and the mean absolute error between the estimated age and chronological age was 3.79 years in the test dataset. In the sex estimation task, our model outputs the probability of male sex (value range from 0 to 1). The overall accuracy was 0.979 (95% confidence interval (CI, 0.967–0.990) and the area under the ROC curve (AUC score) was 0.9975 (95% CI, 0.995–1.000) in the holdout test dataset (**Figs. 2a-c**).

**Table 1.** Multivariate Cox proportional hazards model for primary endpoint.

| Variable (unit) | Coefficient | HR | Confidence interval |  | Z score | P value |
| --- | --- | --- | --- | --- | --- | --- |
|  |  |  | lower 95% | upper 95% |  |  |
| Age (years) | 0.040696 | 1.0415 | 1.0296 | 1.0536 | 6.90978 | $4.85 \times 10^{-12}$ |
| Sex (Male) | 0.076896 | 1.0799 | 0.8992 | 1.2970 | 0.82286 | $4.11 \times 10^{-1}$ |
| BMI (kg/m<sup>2</sup>) | -0.027156 | 0.9732 | 0.9503 | 0.9966 | -2.23923 | $2.51 \times 10^{-2}$ |
| LVEF (%) | -0.015269 | 0.9848 | 0.9782 | 0.9915 | -4.44650 | $8.73 \times 10^{-6}$ |
| log(NT-proBNP) (pg/ml) | -0.117699 | 0.8890 | 0.7180 | 1.1006 | -1.08006 | $2.80 \times 10^{-1}$ |
| Hb (g/dl) | -0.105731 | 0.8997 | 0.8601 | 0.9410 | -4.61192 | $3.99 \times 10^{-6}$ |
| eGFR (ml·min<sup>-1</sup>·1.73 m<sup>-2</sup>) | -0.013147 | 0.9869 | 0.9819 | 0.9920 | -5.02546 | $5.02 \times 10^{-7}$ |
| Age discrepancy (years) | 0.018409 | 1.0186 | 1.0051 | 1.0322 | 2.71196 | $6.69 \times 10^{-3}$ |

Coefficients of the Cox proportional hazards model for the primary endpoint in heart
failure patients. HR, hazard ratio; BMI, body mass index; LVEF, left ventricular
ejection fraction; Hb, hemoglobin; eGFR, estimated glomerular filtration rate; Age
discrepancy, difference between the CXR age and actual age (CXR-age – actual age).

**Fig. 2.**
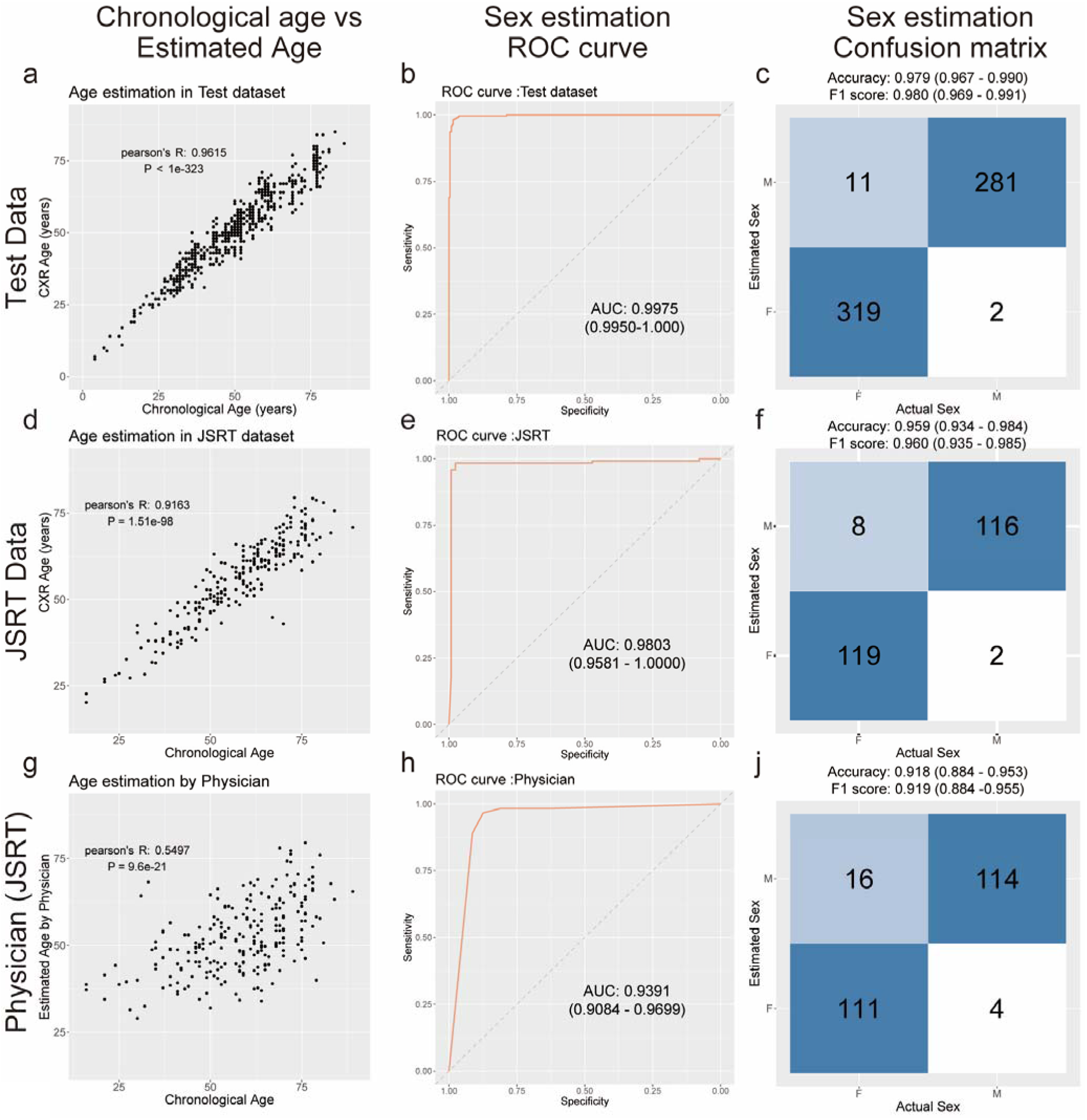
Estimation accuracy of deep learning model and human physician. Estimation accuracy of trained deep learning model in the test dataset (**a**, **b**, **c**), JSRT dataset (**d**, **e**, **f**), and estimation accuracy of human physician in the JSRT dataset (**g**, **h**, **i**). **a**, **d**, **g**, Scatter plots of the actual age (x-axis) and estimated age (y-axis) with Pearson’s correlation coefficient. A strong positive correlation between the actual and estimated age is observed in the deep learning model. In human estimation, the estimated age is the average of the estimations by the four physicians. The correlation between the actual and estimated age was modest (**g**). **b**, **e**, **h**, ROC curves for discriminating between male and female CXRs. The deep learning model accurately estimated sex from CXR. The area under the ROC curve (AUC) and 95% confidence interval are provided. **c**, **f**, **i**, Confusion matrix for sex classification. For the human estimation results, we adopted the average of the estimations by the four physicians (see Methods). Accuracy and F1 metrics and their 95% confidence intervals are displayed at the top.

An important phenomenon known as domain shift sometimes occurs in machine learning, which makes generalization of the machine learning model to unseen data with different distributions difficult ^33^. The NIH chest X-ray data was collected from hospitals in the United States ^25^, most of the patients are likely to be American. To determine whether our model trained using this data can be applied to other populations with different physiques and from different datasets, we also tested the model on the JRST dataset, which is a frontal CXR image dataset comprising 247 frontal CXR images from Japanese patients (**Supplementary Figs. 1b, e**) ^34^. In the JSTR dataset, we also observed a strong significant correlation between the estimated age and chronological age (Pearson’s r: 0.916, p = 1.51 × 10^-98^), and the mean absolute error between the estimated age and chronological age was 4.56 years. Our model also showed high predictive performance in sex estimation (overall accuracy: 0.959 [95% CI, 0.934–0.984] and AUC: 0.9803 [95% CI, 0.9580–1.000]) (**Figs. 2d-f)**. We further examined the reproducibility of this model by extracting images taken multiple times for the same patient from the NIH data. The concordance rate for sex estimation between the two CXRs was 0.982, and for age, the correlation coefficient between the two estimated ages was 0.967 (p < 2.2 × 10^-323^), indicating that both models also showed high reproducibility (**Supplementary Fig. 4**). These results suggest that our model can accurately estimate age and sex from CXR, even in different population groups and cohorts.

### Comparison of predictive performance with human experts

We compared the predictive performance of our model with those of four experienced physicians. We used the JSRT dataset for the comparison because they are familiar with CXR images of Japanese patients. We found a slight correlation between the physicians’ estimated age and actual age, and the average Pearson’s correlation coefficient was 0.38. The mean accuracy and F1 measure in the physicians’ sex estimation were 0.879 and 0.871, respectively. The ensemble predictions by the physicians improved the predictive performance in both age and sex estimations (age: Pearson’s r 0.550, P = 9.6 × 10^-21^, sex: accuracy 0.918 [95% CI, 0.884–0.953], F1 measure 0.919 [95% CI, 0.884–0.955]); however, this did not match the performance of our DNN, particularly in age estimation (**Figs. 2g-i, Supplementary Table 4)**. These results demonstrate that our DNN can learn patterns that are difficult for human experts to recognize.

### Interpretation of deep learning model by heatmap analysis

We attempted to visualize the DNN to understand which part of the image it focused on while estimating the patients’ age and sex. For this purpose, we created a heatmap using Grad-CAM ^35^ and guided backpropagation ^36^. Some examples of images that accurately predicted the patients’ age and sex are shown in **Fig. 3**. For sex classification, the model focused on the breast and clavicle at the upper part of the CXR. This is consistent with the fact that men and women have different amounts of fatty tissue in their breasts. For age estimation, the model mainly focused on the top of the mediastinum and periphery of the rib cage, where the shape and calcification of the aorta seemed to affect the estimation.

**Fig. 3.**
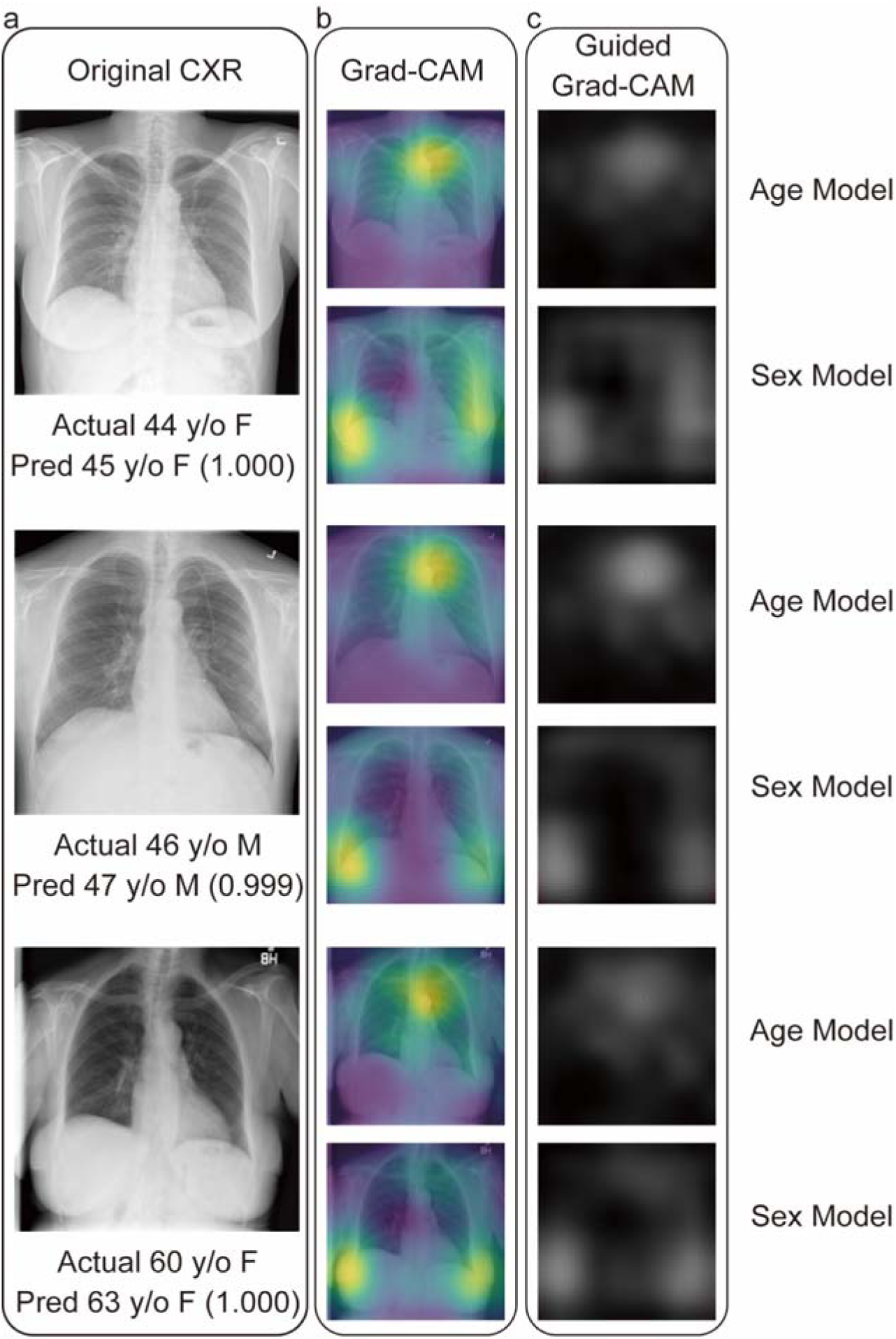
Visualization of the deep learning model with Grad-CAM and guided back-propagation. Example of original CXRs and heatmap visualization using Grad-CAM and guided Grad-CAM. **a**, Original CXR image in the dataset with the actual age, sex, and estimated age and sex (estimated probability). Pred, prediction; F, female; M, male; y/o, years old. **b**, Visualization of the deep learning model using Grad-CAM. Age model (upper row) and sex model (lower row). **c**, Visualization of the deep learning model using a combination of guided backpropagation and Grad-CAM Age model (upper row) and sex model (lower row).

### The difference between the estimated and actual age indicates the existence of a disease

We analyzed CXR images in which the difference between the estimated age and the actual age was large, or sex was incorrectly estimated. Some examples of incorrectly estimated CXRs are shown in **Figs. 4b, c**. Compared with the correctly estimated CXRs (**Fig. 4a**), most of the CXRs with incorrect sex estimation were from children, indicating that sex is difficult to determine in pediatric CXRs. For age estimation, CXRs with a large deviation of estimated age from chronological age seemed to have abnormal findings. For sex prediction, when the performance was evaluated using only the CXR of patients over 20 years old, the accuracy improved from 97.9% to 99.2%. Similarly, for age prediction, the Pearson’s r improved from 0.961 to 0.965 and the mean absolute error improved from 3.79 to 3.66 years in the test dataset when only images with no finding labels were used.

**Fig. 4.**
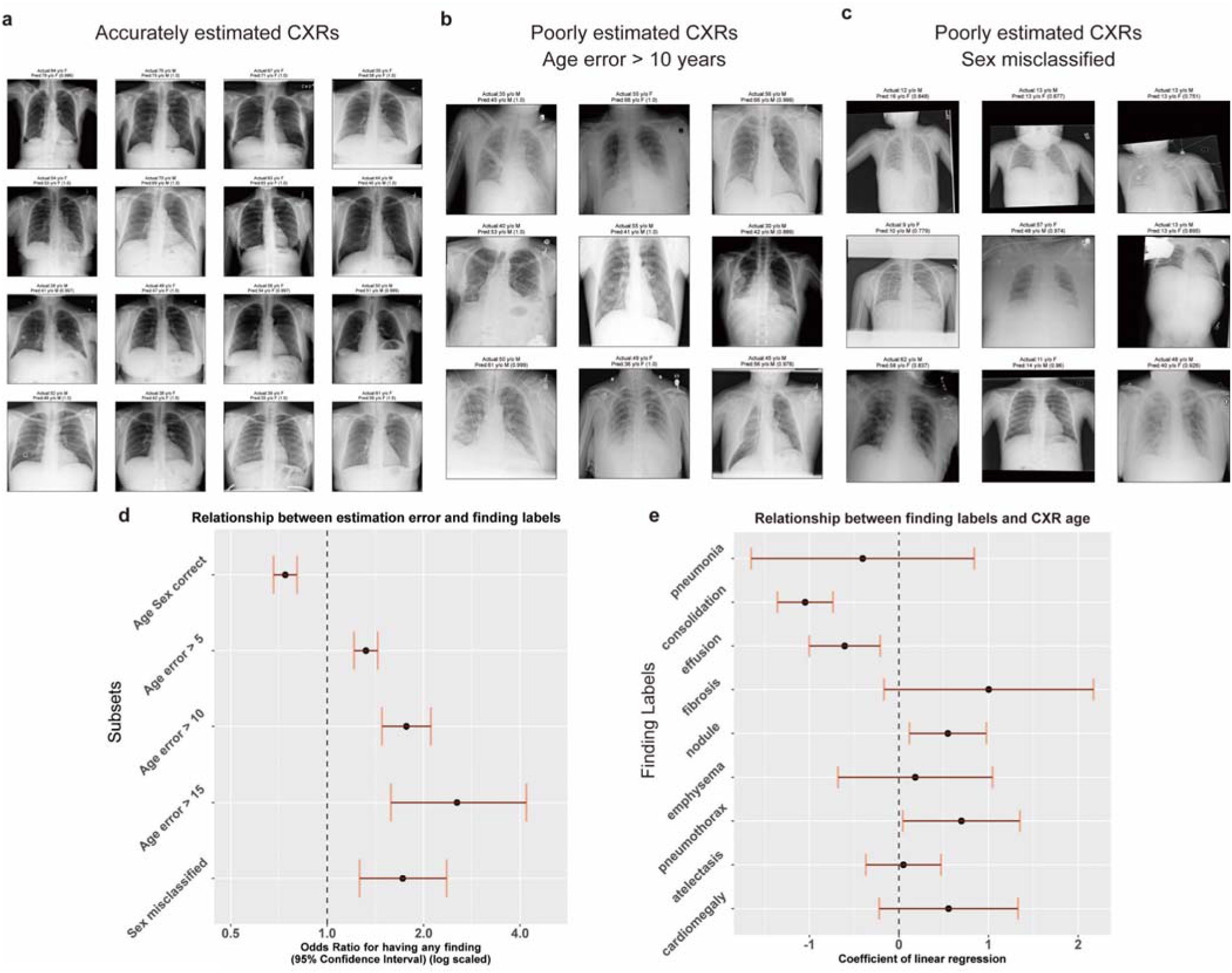
Characteristics of images in which the deep learning model performed inaccurate age and sex estimation. **a**, Example of a CXR image with an age estimation error of less than 5 years and an accurate gender estimation. The actual age, sex, and CXR age and sex (estimated probability) are shown above each image. Pred, prediction; F, female; M, male; y/o, years old. **b**, Example of a CXR image with an age estimation error of more than 10 years. **c**, Example of a CXR image in which the deep learning model failed to estimate its sex correctly. **d**, Relationship between estimation and having any finding labels. The odds ratio with 95% confidence interval is shown on the x-axis. The odds ratio of having any finding labels is lower in CXR images in which the deep learning model correctly estimates their age and sex. On the other hand, images for which gender and age could not be accurately estimated were significantly more likely to have finding labels. **e**, Different finding labels that affect the patient’s estimated age. The coefficient of linear regression adjusted for age (see Methods) is shown on the x-axis.

A large difference was observed between the estimated age and actual age when the images had some finding labels. Conversely, we hypothesize that images with a large deviation of estimated age from the actual age have a higher probability of having some finding labels. We found that CXRs with incorrect sex estimation or a large difference between the estimated and actual ages were significantly more likely to have some finding labels, and this tendency increased with the difference in age (**Fig. 4d**). With respect to each finding label, CXRs with findings of lung nodules and pneumothorax were estimated to be significantly older than the actual age. CXRs with consolidation and effusion yielded the opposite result (**Fig. 4e**). These results suggest that the difference between the estimated and actual age and sex could be a marker for CXR findings, indicating the existence of a disease.

### Estimated age from CXR (CXR age) indicates the presence of cardiovascular abnormalities

The disadvantage of public datasets is that although medical image and imaging findings data are available, they provide little information about the patients. This makes it difficult to investigate a patient’s history and prognosis using public data alone. To address this problem, we used a private database of patients with acute heart failure (HF). This prospective HF registry has enrolled all patients hospitalized for HF since 2011. The registry was designed to collect the clinical background and outcome data of consecutive patients admitted to the Sakakibara Heart Institute for acute decompensated HF. Conventional clinical parameters including age, sex, etiology of HF, risk factors for cardiovascular disease, blood pressure, heart rate, laboratory data, and echocardiographic findings were collected from all study participants (n = 1,562). Events of HF re-hospitalization and death were also recorded ^37–39^. The data comprises 920 (59%) male in the age range of 18–98 years, with a median age of 78 years (interquartile range 69–84) (**Supplementary Figs. 1 c, f**, **Supplementary Table 1**). We applied our model to the CXRs of these patients to estimate their sex and age. The accuracy of sex estimation was 0.945, and the AUC score was 0.986 (95% CI, 0.981– 0.991). Although the performance of age estimation was expected to decrease because all the CXRs were of HF patients and accordingly had some abnormal findings, there was still a significant positive correlation between the estimated and actual age (Pearson’s r: 0.769, p = 4.6 × 10^-291^). We first examined the association between the patient’s history and estimated age from CXR (CXR age) and found that hypertension and atrial fibrillation were significantly associated with increased CXR age after adjustment for chronological age (**Fig. 5a**). Regarding clinical parameters, increased left atrial diameter on echocardiography, tachycardia, and elevated diastolic blood pressure were associated with increased CXR age, whereas increased weight and taller stature were associated with decreased CXR age **(Fig. 5b**). These significant associations suggest that CXR age can be an indicator of cardiovascular abnormalities.

**Fig. 5.**
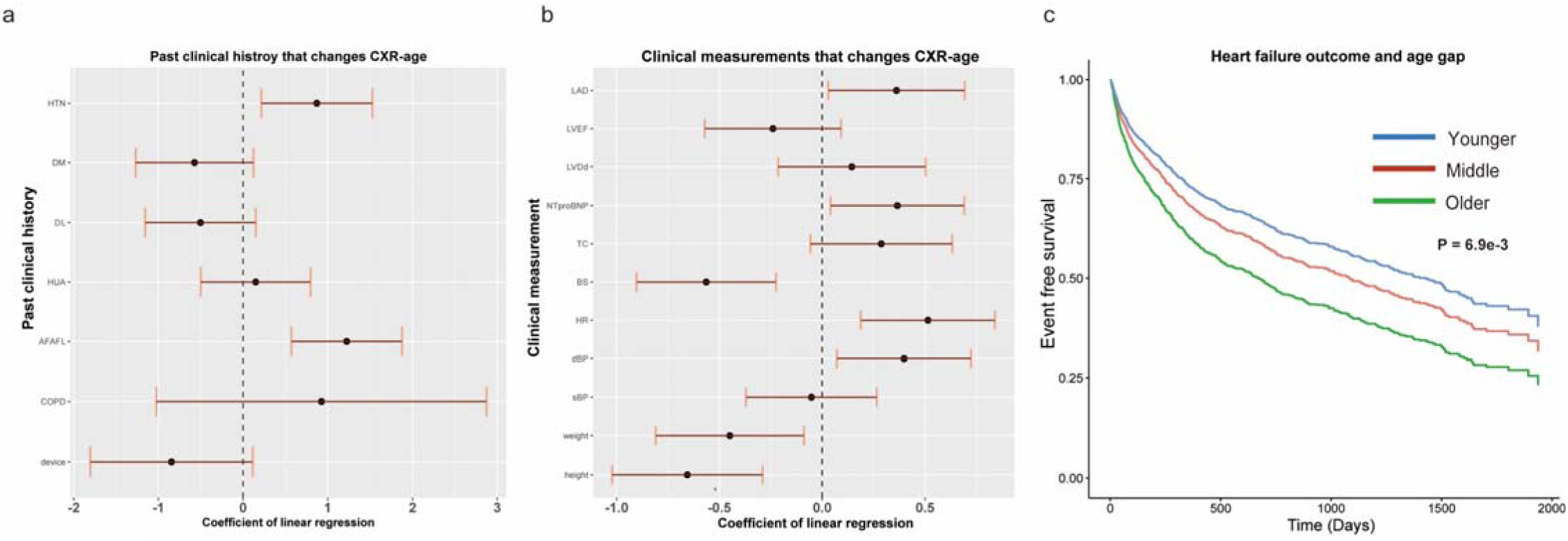
Relationship between CXR age and clinical characteristics and outcome in heart failure patients. Past clinical history (**a**) and continuous clinical measurements (**b**) that affect CXR-age. The coefficient of linear regression adjusted for age is shown on the x-axis with a 95% confidence interval. HTN, hypertension; DM, diabetes mellitus; DL, dyslipidemia; HUA, hyperuricemia; AFAFL, atrial fibrillation or atrial flutter; COPD, chronic obstructive pulmonary disease; device, cardiac pacemaker, implantable cardioverter defibrillator, or cardiac resynchronization therapy devices. LAD, left atrial diameter; LVEF, left ventricular ejection fraction; LVDd, left ventricular end-diastolic diameter; TC, total cholesterol; BS, blood sugar (glucose); HR, heart rate; dBP, diastolic blood pressure; sBP, systolic blood pressure**. c**, Event-free survival curve for heart failure patients stratified by the age discrepancy between the actual and CXR age. Event is defined as the composite endpoint of heart failure re-hospitalization, heart transplantation, and all-cause mortality. The top 20% of patients, middle 60%, and bottom 20% were grouped as older, middle, and younger, respectively.

### CXR age predicts heart failure prognosis

Next, we examined the association between HF outcomes and CXR age. We defined the primary endpoint as the composite endpoint of all-cause mortality and HF re-hospitalization. In the univariate Cox proportional hazards model, CXR age is associated with the primary endpoint as well as other conventional risk factors such as age, sex, body mass index (BMI), hemoglobin (Hb), NT-pro BNP, and eGFR (**Supplementary Table 3**). Sex misclassification was positively associated with worse outcomes, but not significantly. For multivariate analysis, the difference between CXR age and chronological age was independently associated with the primary endpoint after adjustment for conventional risk factors, suggesting that patients estimated to be older had a worse HF prognosis (**Fig. 5c, Table 1**). The Akaike information criterion (AIC) and Bayesian information criterion (BIC) are often used as a criterion for better model selection, and lower values suggest a better model for this criterion. Interestingly, AIC and BIC were decreased in the Cox model by replacing the age of the conventional model with CXR age (**Supplementary Table 6**), indicating that CXR age can be a better prognostic indicator than actual age.

## Discussion

In this study, we verified the performance of a DNN to estimate patients’ age and sex from CXRs without any additional clinical data. We also explored the clinical implications of the estimated age and sex. The main findings of the present study are summarized below. 1) The patient’s age was estimated from CXR within 5 years of mean absolute error using a deep learning algorithm. The patient’s sex was also estimated from CXR with more than 95% accuracy. 2) Our DNN estimations of age and sex were much more accurate than the ensemble estimation made by cardiovascular and respiratory medicine experts. 3) Our model focused on the breast and the area around clavicle for sex estimation and on the mediastinum for age estimation. 4) The difference between CXR age and actual age was large in CXRs with abnormal findings. 5) In the HF population, patients with hypertension and atrial fibrillation were estimated to be older. CXR age was independently associated with HF outcomes after adjusting for covariates. From these findings, we conclude that age and sex can be estimated from CXR with high accuracy and reproducibility using our model, and that CXR age can be used as a simple measure of health status in patients with cardiovascular disease.

Although age and sex affect CXR findings, few previous studies have reported age and sex estimation from CXR images ^12, 13^. Karargyris et al. reported a CNN model that predicts age from CXR using the NIH dataset. However, they only reported the predictive performance on internal validation datasets, which can lead to overestimation because validation data was used for tuning the hyperparameters of the model. The model performance should be evaluated using unseen data ^40, 41^. Our evaluation method is robust and fair in that we evaluated the estimation performance on an external test dataset and entirely independent JSRT dataset, both of which were not used during the training phase. We also visualized our CNN model using Grad-CAM and guided-Grad-CAM and found that our model focused around the top of the mediastinum in predicting the patient’s age. Tortuosity and calcification of the aorta have been reported to be characteristic of atherosclerotic disease ^42–44^. On the other hand, to the best of our knowledge, this is the first study to estimate sex from CXR using deep learning. Our model seemed to focus not only on the breast as expected, but also around the clavicle. The length and shape of the clavicle are reported to be different in males and females ^45, 46^. Our heatmap analysis results were consistent with those reported in these previous reports.

Our DNN can be applied in various ways. For instance, the sex estimation model exhibited high accuracy and could be used as an annotation tool for anonymous medical data or could be employed to generate an alert to prevent patient mix-ups in clinical practice. The age estimation model can provide a simple biomarker that represents a single quantification of information from the entire CXR image. A discrepancy between CXR age and chronological age suggests the presence of abnormal findings in the CXR image. Actually we found that patients with older CXR age had a significantly higher probability of hypertension and atrial fibrillation, both of which are related to cardiovascular aging ^47, 48^. CXR age was also associated with worse outcomes in patients with HF. Our results suggest that CXR age can be a simple health indicator that reflects the aging state of the heart and vessels. As an indicator of the degree of aging, perceived age is a robust biomarker that has been linked to age-related diseases and prognosis. However, age estimation by a single physician is not an objective indicator ^2–4^. Furthermore, it is not easy to take facial photographs of patients in clinical settings due to privacy concerns, which hinders the clinical application of perceived age. Since CXR is taken in most patients as a screening test, the estimation of aging by CXR age has the potential to replace perceived age as an objective biomarker. There are several methods to determine a patient’s health status from laboratory data based on age. For example, “lung age” estimated from spirometry forced expiratory volume (FEV) ^49^ and “vascular age” estimated from carotid artery ultrasonography ^50^ are used as simple health indicators in clinical practice, and these methods help clinicians explain test results to patients. With continued advancement in deep learning as demonstrated in this study of CXR age, medical images will also be quantified as an age. Additionally, our study shows it possible to digitize CXR images into a single numerical value, which presents a new possibility. Namely, by quantifying CXR images, we can successfully incorporate them as parameters in clinical studies, which has been difficult in the past. For instance, in clinical studies of HF or cardiovascular diseases, laboratory data, echocardiographic parameters (such as left ventricular ejection fraction or left atrial dimension), and ECG arrhythmia categories are included in statistical variables as these are numerical values; however, CXR image findings are difficult to include as variables. Although there is an index of cardiothoracic ratio, it is difficult to quantify the entire CXR information. This method can be applied to other medical images, such as CT, MRI, and ultrasonography. Our results demonstrate the potential for new applications of CXRs, the most widely performed imaging test in the world.

The high accuracy of deep learning has been reported in the diagnosis analysis of various medical images such as skin images, pathology slides, ECGs, CXRs, CT, MRI, and echocardiography^15, 16, 19, 20, 51–54^. Several studies reported that deep learning can accomplish tasks that are even difficult for human physicians ^21, 22, 55^. In the example of CXR, Lu et al. created a deep learning model to predict mortality risk from CXR images and stratified the risk of long-term mortality ^56^. Toba et al. reported that they estimated the pulmonary to systemic flow ratio, an indicator of the severity of congenital heart disease, from CXR ^57^. However, few studies have reported age estimation using medical imaging. It has been reported that age estimation from hand and knee MRI using deep learning can estimate the age of young people with high accuracy ^58, 59^. Attia et al. created a deep learning model to predict age and sex from a 12-lead ECG and achieved a classification accuracy of 90.4% for sex estimation and an MAE of 6.9 years for age estimation. They also reported that patients with a predicted age exceeding the true age by more than 7 years had a higher incidence of lower cardiac function, hypertension, and coronary artery disease ^23^. Wang et al. proposed a deep learning model to predict patients’ age from brain MRI and reported that the estimated age is associated with dementia ^24^. Although these previous studies have reported that image-estimated age is associated with disease, our study is the first to identify its association with disease prognosis.

The present study also has several limitations. First, all CXR images were obtained from patients, and hence, they were obtained for some clinical indications. Therefore, it is unclear whether our results would be applicable to healthy population. We only examined the relationship between estimated age, disease, and prognosis in patients with heart failure. Further studies are needed to determine if this is applicable to other patients to the general population; for instance, using a large amount of data from medical checkups. Second, as is often the case with large datasets, the NIH chest X-ray dataset contains low-quality images and labels. Finding labels may not necessarily be accurate because the NIH dataset was labeled using natural language processing ^25^. Third, the analysis of our model heart failure is a single-center observational study with a relatively modest number of patients, and the findings of the study can potentially include some bias due to its retrospective nature. Fourth, as described above, older estimated age does not necessarily mean worse CXR findings. For instance, CXRs with findings of consolidation or effusion were estimated to be younger than the actual age. However, analysis of a large dataset and data for heart failure patients suggests that CXR age reflects, to some extent, aging and health status.

In conclusion, we developed DNNs that estimates patients’ age and sex from CXR without any additional information, and our models exhibited a high predictive performance with the independent JSRT dataset. Our study suggests that CXR age can serve as a novel biomarker for cardiovascular aging and health status, and can be a key tool to help clinicians predict, prevent, and manage cardiovascular diseases in the era of digital medicine.

## Methods

### Dataset acquisition

Three datasets were used in this study (**Fig. 1 and Supplementary Fig. 1**). We used the NIH ChestX-ray, which comprises 112,120 png images of frontal-view CXR from 30,805 unique patients. This dataset also includes metadata containing patients’ age and sex information with up to 15 labels ^25^. We excluded 16 CXR images from patients over 100 years of age. We randomly split the dataset into three groups (training set: 102,029 images from 28,029 patients; validation set: 9,426 images from 2,523 patients; test sets: 613 images from 250 patients). There was no patient overlap between the sets to avoid data leakage during model training, which can lead to overestimation of model performance. Although deep learning models were trained separately for age and sex estimation, we used the same data split for training, validation, and test sets for both tasks. We also used the JSRT database, which comprises 247 frontal CXR images from Japanese patients with or without lung nodules^34^. We removed two images for which age information was not available. The JSRT database was used as an independent test dataset to check the generalizability of our model and to determine whether our model can be applied to other populations with different physiques. Heart failure patient data were obtained from our prospective heart failure registry, which enrolled all patients with acute decompensated heart failure who were admitted to Sakakibara Heart Institute (Fuchu, Tokyo), a hospital specializing in cardiovascular disease. The diagnosis of heart failure was based on the Framingham criteria ^60^. Patients with acute coronary syndrome and isolated right-sided HF were excluded. Conventional clinical variables including age, sex, etiology of HF, risk factors, blood pressure, heart rate, laboratory data, and echocardiographic findings were obtained from all the study participants. Events of heart failure, re-hospitalization, and death were also recorded. Frontal CXRs within 2 days of hospital admission were used in the analysis. Written informed consent was obtained from all the participants before the study. The study protocol was also approved by the Institutional Review Board of Sakakibara Heart Institute (No. 19-092).

### Deep learning model development and training

To develop a deep learning model for age and sex estimation, we applied transfer learning and fine-tuning techniques to our model. We adopted 11 convolutional neural network (CNN) architectures, namely ResNet18, ResNet34, ResNet50, ResNet101, ResNet152 ^28^, DenseNet121, DenseNet161, DenseNet169, DenseNet201 ^29^, Inception-v4 ^30^ and SENet154 ^31^. For transfer learning, we used pre-trained weights for the CNN models. Pre-trained weights on ImageNet were downloaded for each model from https://github.com/Cadene/pretrained-models.pytorch. Models can be separated into two parts in a CNN: the convolutional part and fully connected layer (FCL) part. Because these models are for the classification task of 1000 categories, the default output layer is comprised of 1000 neurons, which represent the probabilities of each category (**Fig. 1**). The convolutional layers were initialized with loaded pre-trained weights and were frozen. We modified the original FCL part into a new two-layered FCL part. The FCL part is composed of batch-normalization, an FCL of 512 neurons with a rectified linear unit (ReLU) as the activation function, batch normalization ^61^ and a final FCL. The final layer neuron outputs the probabilities of males and females for the sex estimation task. Dropout ^62^ was applied after batch normalization. For age estimation, we adopted an FCL with a single final neuron so that the model outputs a single numerical value of the predicted age and makes it a regression problem. We selected (1) binary cross-entropy (BCE) loss and (2) mean square error (MSE) loss for sex and age estimation, respectively. BCE and MSE are defined by the following equations, where **n** is the number of images, *p*_male_ is the estimated probability of male sex, **y** is the actual label, and 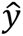 is the estimated age.

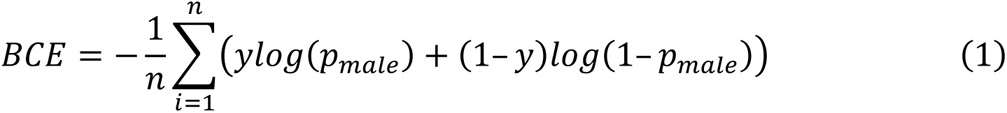

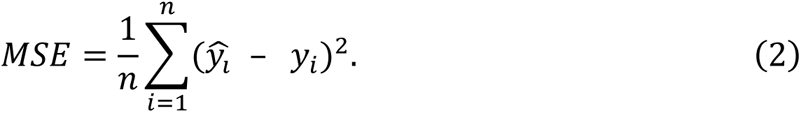

The models were trained on the training dataset to minimize these loss functions. The models were trained using the Adam optimizer and a cyclic learning rate policy ^63^. During transfer learning, only the parameters in the FCL part and the batch norm layer of the convolutional part were updated. Then, we fine-tuned the entire network by unfreezing and updating the pre-trained weights with a much lower learning rate. The validation set was used to select hyperparameters to determine when to stop training to avoid overfitting and to select the final model. Validation data was not used to update the weights of the DNN model. The NIH ChestX-ray database provides png images, and the JSRT and heart failure patients’ CXRs were DICOM images. All the images were transformed into png images using Python’s ‘pydicom’ library and resized to 320 × 320 pixels. To improve the generalizability of our model and avoid overfitting, we applied image augmentation ^32^. The images in the training datasets were augmented with random padding and random rotation up to ± 20°. Image flipping was not performed. Our DNNs were trained on NVIDIA Tesla V100 GPUs with a mixed precision training technique ^64^. After the training, we selected the model with the lowest loss value in the validation dataset as the final model (**Supplementary Tables 2 and 3**). We applied the trained DNN to the test dataset and JSRT dataset to assess the estimation performance. Image augmentation was not applied to the test and JSRT datasets. We used gradient-weighted class activation mapping (Grad-CAM) ^35^ and guided backpropagation ^36^ methods to visualize the area of interest of our models.

### Age and sex estimation by human physicians

To compare our model with human physicians’ prediction performance, four trained clinicians (three cardiologists and one pulmonologist) estimated the patient’s age and sex from CXRs on the JSRT dataset. They have 8, 9, 15, and 30+ years of clinical experience, respectively. They estimated the patient’s age from the CXR image without any additional information. We used the JSRT data because they are physicians in Japan and are usually accustomed to diagnosing Japanese CXRs. They were allowed to see the training dataset images and labels before estimating age and sex in the JSRT dataset. For ensemble prediction, the estimated age and sex of the four physicians were averaged. For instance, ensemble prediction is 40-year-old and 0.75 probability for male when the four physicians estimate a CXR as a 48-year-old male, 52-year-old male, 55-year-old male, and 25-year-old female.

### Statistical analysis of test results

To estimate the predictive performance of the model, the Pearson’s R value for age estimation was calculated. The mean squared error and mean absolute error were calculated. For the sex model, we computed the area under the receiver operating characteristic curve (AUC) to assess the sex model discrimination performance. The classification accuracy and F1 score were also calculated. The confidence intervals for AUC, accuracy, and F1 metrics were derived from the 100,000 bootstrap replications method. To test the model’s reproducibility, we extracted patients who had multiple CXRs within one year in the validation and test data. Age and sex were estimated from CXR using our DNN, and the Pearson’s r correlation coefficient was calculated for age and the concordance rate for sex estimation was calculated. Linear regression was used to analyze the association between estimated age and finding labels, and the coefficient of disease label was determined. The three finding labels of edema, infiltration, and consolidation were grouped together as consolidation, and hernia was excluded from the analysis because it was labeled in a small number of CXR images (227 images out of 112,104 images). The Cox proportional hazards model was used for survival analysis. The median follow-up period was 407 days (interquartile range: 122–879). Event was defined as the composite endpoint of heart failure re-hospitalization and all-cause mortality. The independent variables in the Cox model were determined by referring to empirical rules and previous articles. Age, sex, BMI, previous history of hypertension, diabetes mellitus, dyslipidemia and smoking, LVEF, NT-pro BNP, Hb, eGFR, CXR age, and sex misclassification by the deep learning model were incorporated as independent variables. Variable selection for the multivariate analysis, age, sex, and left ventricular ejection fraction (LVEF) were fixed as independent variables because they are known to be strong predictors of heart failure outcome ^65, 66^. Independent variables that showed P values of less than 0.05 in univariate analysis were employed in the multivariate analysis. To compare the Cox model and different independent variables, we used the AIC and BIC. The AIC and BIC values were tested using the 100,000 bootstrap replications method. The R version 3.6.3 base function and ‘caret’, ‘survival’, and ‘boot’ packages were used for all statistical analyses. A raw two-sided p value is provided when the p value is greater than 2.2 × 10^-323^, otherwise it is provided as p < 2.2 × 10^-323^ because of generic computational limitations.

### Data availability

The dataset generated and analyzed during this study is available from the corresponding authors on request. The NIH ChestX-ray dataset used in this study is openly available and can be downloaded at https://cloud.google.com/healthcare/docs/resources/public-datasets/nih-chest. The JSRT database used in this study is also publicly available and can be downloaded at http://db.jsrt.or.jp/eng.php.

### Code availability

The code is available on request.

## Acknowledgements

We would like to express our gratitude to the members of the Sakakibara Heart Institute for their support in collecting samples and providing clinical information. We are grateful to the National Institute of Health and the Japanese Society of Radiological Technology for making their data publicly available.

## Author Contributions

H.I., K.I., M.S., Y.N, and T.Y. conceived and designed the study. M.S, Y.N., and T.Y. collected and managed the heart failure patient sample. H.I and K.I. developed the deep learning model and performed statistical analysis. R.K. provided computer resources and intellectual advice for developing the deep learning model. Y.N., S. Koyama, H.M., K.M., K.O., Y.O., S. Katsushika, R.M., H.S., T.Y., S. Kodera, Y.H., K.F., and H.A. contributed to data analysis and interpretation. M.I., T.Y., and I.K. supervised the study. H.I. and K.I. wrote the manuscript, and many authors also provided valuable edits.

