## Supplementary Materials for "Deep learning-based chest X-ray age serves as a novel biomarker for cardiovascular aging"

### Supplementary Tables

##### Supplementary Table 1. Characteristics of heart failure patients

| Clinical measurements (unit) | Heart failure patients (n=1562) |
|---|---|
| Age (years old) | 78 [69, 84] |
| Sex (Male) | 920 (58.9) |
| Height (cm) | 159 [151.5, 166.5] |
| Weight (kg) | 58.3 [50, 67.8] |
| BMI ($kg/m^2$) | 23.1 [20.7, 25.8] |
| Etiology of heart disease | |
| ischemic | 424 (27.1) |
| valvular | 537 (34.4) |
| other | 601 (38.5) |
| Hypertension | 945 (60.5) |
| Diabetes mellitus | 446 (28.6) |
| Dyslipidemia | 625 (40.0) |
| Smoking history | 760 (48.7) |
| Atrial fibrillation/atrial flutter | 974 (62.4) |
| HOT | 40 (2.6) |
| Implantable device | |
| None | 1359 (87.0) |
| PM | 127 (8.1) |
| ICD | 53 (3.4) |
| CRT | 23 (1.5) |
| Systolic BP (mmHg) | 137 [119, 157] |
| Diastolic BP (mmHg) | 79 [66, 95] |
| Heart rate (min) | 89 [71, 110] |
| SpO2 (%) | 95 [92, 98] |
| Clinical scenario | |
| CS1 | 728 (46.6) |
| CS2 | 719 (46.0) |

| | |
|---|---|
| CS3 | 115 (7.4) |
| Laboratory measurements | |
| Hb (g/dl) | 12.0 [10.5, 13.6] |
| Hct (%) | 37.0 [32.6, 42.0] |
| BUN (mg/dl) | 21.2 [16.5, 29.4] |
| Na (mEq/L) | 140 [137, 142] |
| K (mEq/L) | 4.4 [4.0, 4.7] |
| T- Bil (mg/dl) | 1.0 [0.7, 1.4] |
| AST (U/L) | 35 [26, 52] |
| ALT (U/L) | 23 [15, 41] |
| ALP (U/L) | 292.38 (216.32) |
| Albumin (U/L) | 3.7 [3.3, 3.9] |
| UA (U/L) | 6.4 [5.3, 7.8] |
| NT-proBNP (pg/ml) | 3777 [1952, 7653] |
| CRP (mg/dl) | 0.43 [0.14, 1.40] |
| Cre (mg/dl) | 1.00 [0.79, 1.33] |
| eGFR ($ml \cdot min^{-1} \cdot 1.73\ m^{-2}$) | 51.2 [36.47, 64.07] |
| WBC | 6400 [5100, 8200] |
| Lymph (%) | 20.80 [14.70, 28] |
| BS (mg/dl) | 124 [105, 157.5] |
| TSH (µIU/ml) | 2.44 [1.51, 4.21] |
| HbA1c (%) | 5.9 [5.5, 6.4] |
| TC (mg/dl) | 159 [135.75, 185] |
| Echocardiography parameters | |
| LVDd (mm) | 51 [44, 58] |
| LVDs (mm) | 38 [31, 49] |
| LVEF (%) | 47 [31, 58] |
| LAD (mm) | 45 [40, 51] |
| TRPG (mmHg) | 28 [22, 37] |

Characteristics of patients with HF in this study. Continuous variables are presented as median [interquartile range], except for ALP. ALP is expressed as mean (standard deviation), as it was normally distributed, using the Shapiro–Wilk test. Categorical variables are presented as n (%).

BMI, body mass index; HOT, home oxygen therapy; PM, pacemaker; ICD, implantable cardioverter defibrillator; CRT, cardiac resynchronization therapy; Hb, hemoglobin; BUN, blood urea nitrogen; T-Bil, total bilirubin; AST, aspartate aminotransferase; ALT, alanine aminotransferase; ALP, alkaline phosphatase; UA, urinary acid; CRP, C-reactive protein, Cre, creatinine; eGFR, estimated glomerular filtration rate; WBC, white blood cell count; Lymph, lymphocytes; BS, blood glucose; TSH, thyroid stimulating hormone; HbA1c, hemoglobin A1C; TC, total cholesterol; LVDd, left ventricular end-diastolic diameter; LVDs, left ventricular end-systolic diameter; LVEF, left ventricular ejection fraction; LAD, left atrial dimension; TRPG, tricuspid regurgitation peak gradient.

**Supplementary Table 2. Estimation accuracy of age model in the validation dataset using different deep learning architectures.**

| Architecture | MSE | RMSE | R | MAE |
|---|---|---|---|---|
| ResNet18 | 34.96 | 5.912 | 0.9364 | 4.5450 |
| ResNet34 | 32.75 | 5.723 | 0.9399 | 4.4196 |
| ResNet50 | 30.02 | 5.479 | 0.9460 | 4.2026 |
| ResNet101 | 29.84 | 5.463 | 0.9462 | 4.2045 |
| ResNet152 | 32.85 | 5.731 | 0.9423 | 4.2248 |
| DenseNet121 | 30.66 | 5.537 | 0.9449 | 4.2511 |
| DenseNet161 | 30.30 | 5.504 | 0.9447 | 4.2156 |
| DenseNet169 | 33.36 | 5.776 | 0.9390 | 4.2473 |
| DenseNet201 | 27.69 | 5.262 | 0.9493 | 4.0373 |
| Inception v4 | 31.04 | 5.571 | 0.9449 | 4.3174 |
| **SENet154** | **27.34** | 5.229 | 0.9524 | 4.0768 |

The model with the smallest mean squared loss in the validation dataset was selected as the final model. MSE, mean squared error; RMSE, root mean squared error; R, Pearson's r between the actual and estimated age; MAE, mean absolute error.

**Supplementary Table 3. Estimation accuracy of sex estimation model in the validation dataset using different deep learning architectures.**

| Architecture | BCE | Accuracy | AUC |
|---|---|---|---|
| ResNet18 | 0.0568 | 0.9766 | 0.9980 |
| ResNet34 | 0.0527 | 0.9769 | 0.9983 |
| ResNet50 | 0.0489 | 0.9790 | 0.9984 |
| ResNet101 | 0.0530 | 0.9822 | 0.9987 |
| ResNet152 | 0.0533 | 0.9790 | 0.9984 |
| DenseNet121 | 0.0519 | 0.9800 | 0.9983 |
| **DenseNet161** | **0.0430** | 0.9826 | 0.9988 |
| DenseNet169 | 0.0508 | 0.9792 | 0.9985 |
| DenseNet201 | 0.0518 | 0.9802 | 0.9986 |
| Inception v4 | 0.0458 | 0.9819 | 0.9989 |
| SENet154 | 0.0571 | 0.9788 | 0.9981 |

The model with the smallest binary cross-entropy loss in the validation dataset was selected as the final model. BCE, binary cross-entropy loss; AUC, area under the ROC curve.

**Supplementary Table 4. Summary of human physician estimation of age and sex.**

| | R | Accuracy | F1 |
|---|---|---|---|
| Cardiologist 1 | 0.483 | 0.833 | 0.811 |
| Cardiologist 2 | 0.204 | 0.902 | 0.901 |
| Cardiologist 3 | 0.440 | 0.878 | 0.868 |
| Pulmonologist 4 | 0.392 | 0.902 | 0.905 |
| Physician performance mean | 0.380 | 0.879 | 0.871 |
| Physician ensemble prediction | 0.550 | 0.918 | 0.917 |
| DNN | 0.916 | 0.959 | 0.960 |

Age and sex estimation performance of human physicians in the JSRT dataset. The predictive performances of the four physicians are given. The physician performance mean is the average of the four physicians' metrics. The physician ensemble prediction is the ensemble prediction metric (see the Methods section). R, Pearson's r between the actual and estimated age; Accuracy, overall accuracy of sex estimation; F1, f1 score for sex estimation; DNN, deep neural network.

**Supplementary Table 5. Cox proportional hazards model for primary endpoint.**

| Variable (unit) | Coefficient | OR | Confidence interval | | z | P value |
|---|---|---|---|---|---|---|
| | | | lower 95% | upper 95% | | |
| Age (years) | 0.03911 | 1.0399 | 1.0316 | 1.0482 | 9.56636 | $1.11\times10^{-21}$ |
| Sex (Male) | -0.15344 | 0.8578 | 0.7315 | 1.0058 | -1.88893 | $5.89\times10^{-2}$ |
| BMI (kg/m$^2$) | -0.05290 | 0.9485 | 0.9286 | 0.9688 | -4.90035 | $9.57\times10^{-7}$ |
| Hypertension | -0.12996 | 0.8781 | 0.7477 | 1.0313 | -1.58418 | $1.13\times10^{-1}$ |
| Diabetes mellitus | 0.07798 | 1.0811 | 0.9096 | 1.2849 | 0.88495 | $3.76\times10^{-1}$ |
| Dyslipidemia | 0.04725 | 1.0484 | 0.8941 | 1.2293 | 0.58155 | $5.61\times10^{-1}$ |
| Smoking history | -0.04413 | 0.9568 | 0.8171 | 1.1204 | -0.54794 | $5.84\times10^{-1}$ |
| LVEF (%) | -0.00369 | 0.9963 | 0.9912 | 1.0015 | -1.40297 | $1.61\times10^{-1}$ |
| log (NT-proBNP) (pg/ml) | 0.50941 | 1.6643 | 1.4055 | 1.9708 | 5.90755 | $3.47\times10^{-9}$ |
| Hb (g/dl) | -0.15665 | 0.8550 | 0.8244 | 0.8868 | -8.41689 | $3.87\times10^{-17}$ |
| eGFR (ml·min$^{-1}$·1.73 m$^{-2}$) | -0.01864 | 0.9815 | 0.9775 | 0.9855 | -8.97716 | $2.78\times10^{-19}$ |
| CXR age (years) | 0.03963 | 1.0404 | 1.0308 | 1.0502 | 8.31358 | $9.29\times10^{-17}$ |
| Sex misclassification | 0.23516 | 1.2651 | 0.9069 | 1.7648 | 1.38462 | $1.66\times10^{-1}$ |

Coefficients of the univariate Cox proportional hazards model for the primary endpoint in heart failure patients. Hypertension, diabetes mellitus, dyslipidemia, smoking history, and sex misclassification were treated as binary categorical variables. OR, odds ratio; BMI, body mass index; LVEF, left ventricular ejection fraction; Hb, hemoglobin; eGFR, estimated glomerular filtration rate.

**Supplementary Table 6. Comparisons of different Cox proportional hazards models.**

| Model | Covariates | AIC | P value (vs model 1) | BIC | P value (vs model 1) |
|---|---|---|---|---|---|
| Model 1 (Nominal) | Age + Sex + BMI + LVEF + log(NT-proBNP) + Hb + eGFR | 3625.0 | - | 3650.7 | - |
| Model 2 | Age + Sex + BMI + LVEF + log(NT-proBNP) + Hb + eGFR + Age_discrepancy | 3437.4 | $< 2 \times 10^{-5}$ | 3466.4 | $< 2 \times 10^{-5}$ |
| Model 3 | CXR age + Sex + BMI + LVEF + log(NT-proBNP) + Hb + eGFR | 3456.5 | $4.00 \times 10^{-5}$ | 3481.8 | $4.00 \times 10^{-5}$ |

Comparison of different Cox proportional hazards models with different variables. The P value was calculated using the 100,000 bootstrap replications method. BMI, body mass index; LVEF, left ventricular ejection fraction; Hb, hemoglobin; eGFR, estimated glomerular filtration rate; CXR age, age estimated using the deep learning model; Age_discrepancy, difference between CXR age and actual age (CXR age - actual age).

### Supplementary Figures

**Supplementary Fig. 1 Age and sex distribution in the datasets.**

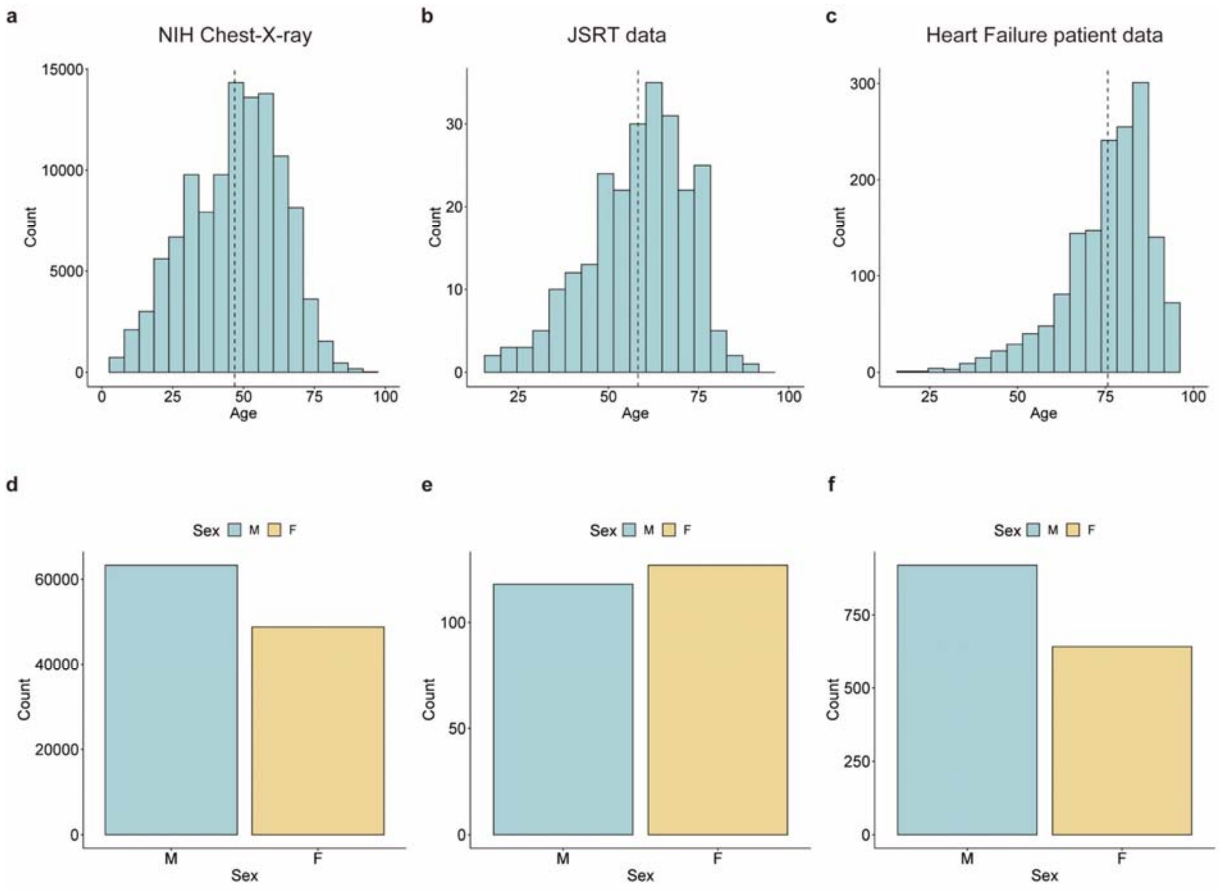


Histogram of age and sex in the NIH Chest-X-ray database (**a**, **d**), JSRT database (**b**, **e**), and heart failure patient data (**c**, **f**). M, male; F, female.

**Supplementary Fig. 2 Study flowchart and data usage.**

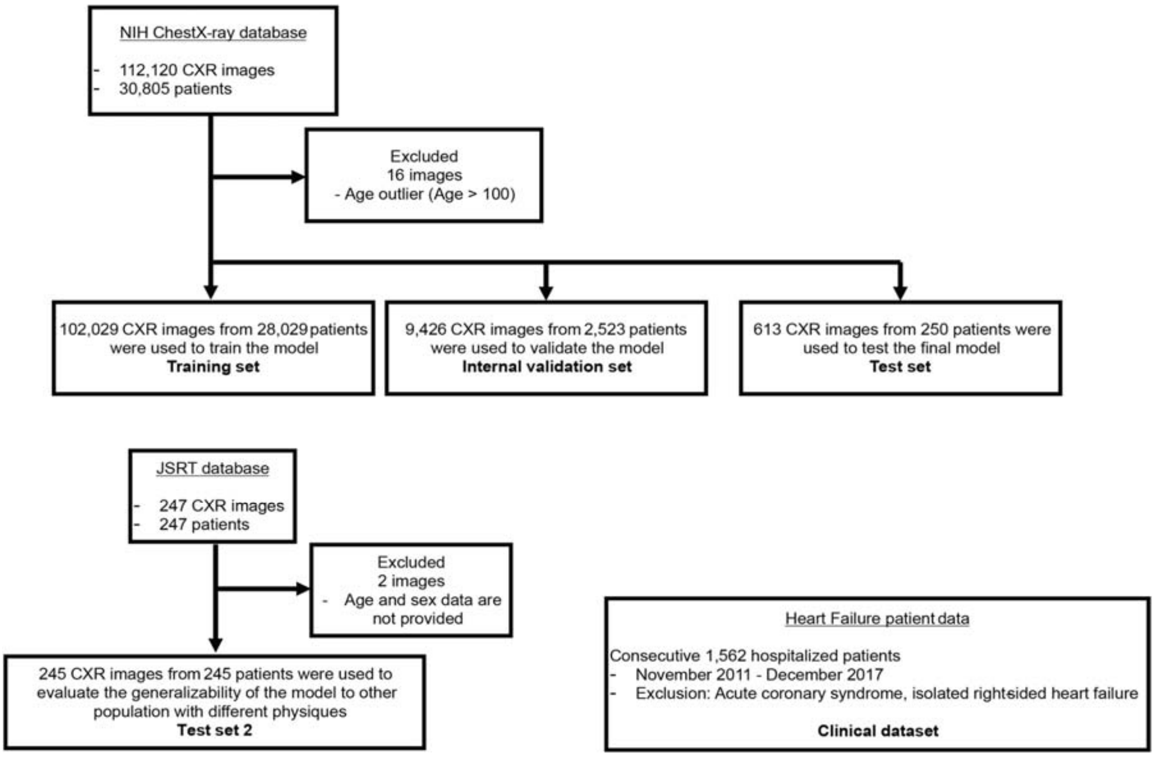


Three datasets were used in this study. The NIH Chest X-ray database was randomly divided into training, validation, and test datasets after exclusion of 16 age outliers. The JSRT database was used as the independent external test data to evaluate the predictive performance of the deep learning model. Two patients whose age and sex data were unavailable were excluded. The heart failure patient data consists of 1,562 consecutive patients who were admitted to Sakakibara Heart Institute and was used to explore the clinical significance of estimated age in real-world patient data.

**Supplementary Fig. 3 Estimation accuracy of the deep learning model in validation data.**

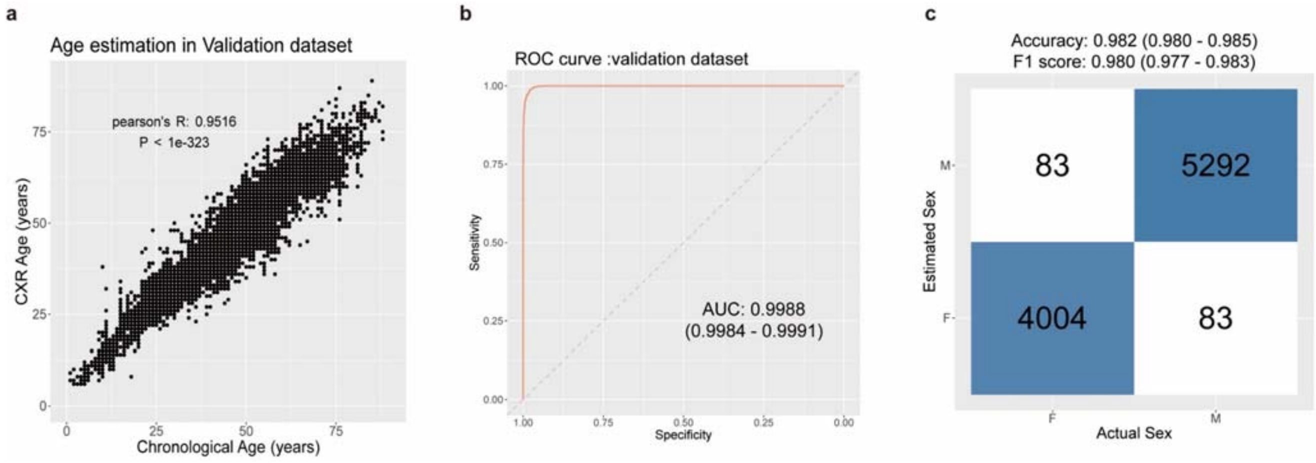


**a**, Scatter plot of the actual age (x-axis) and estimated age (y-axis) with Pearson's correlation coefficient in the validation dataset. **b**, ROC curve for discriminating male and female from CXRs with AUC values and 95% confidence intervals. AUC, area under the ROC curve. **c**, Confusion matrix for sex classification. Accuracy and F1 metrics and their 95% confidence intervals are displayed at the top.

**Supplementary Fig. 4 Reproducibility analysis.**

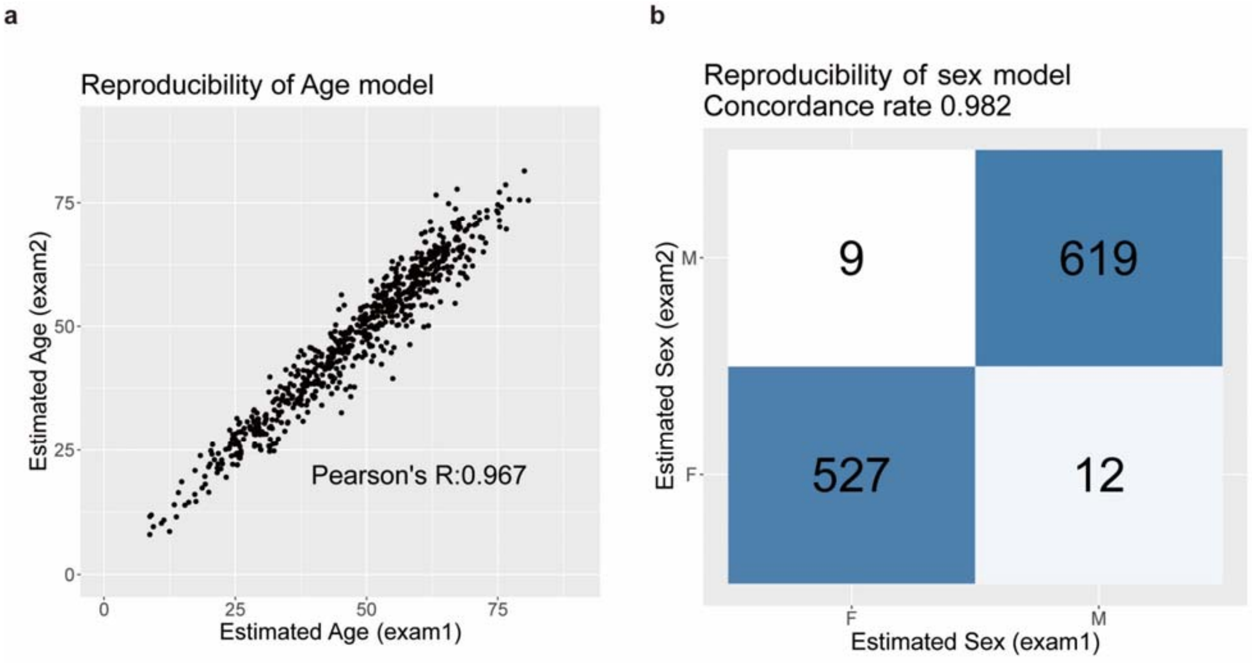


**a** Scatter plot of the estimated age of different CXRs from the same patients. The estimated ages of the first and second exams are plotted on the x- and y-axes, respectively. **b**, Confusion matrix of the estimated sex of different CXRs from the same patients. The estimated sexes of the first and second exams are given on the x- and y-axes, respectively.

**Supplementary Fig. 5 Histogram of bootstrapping statistics of AIC and BIC in the Cox proportional hazards model.**

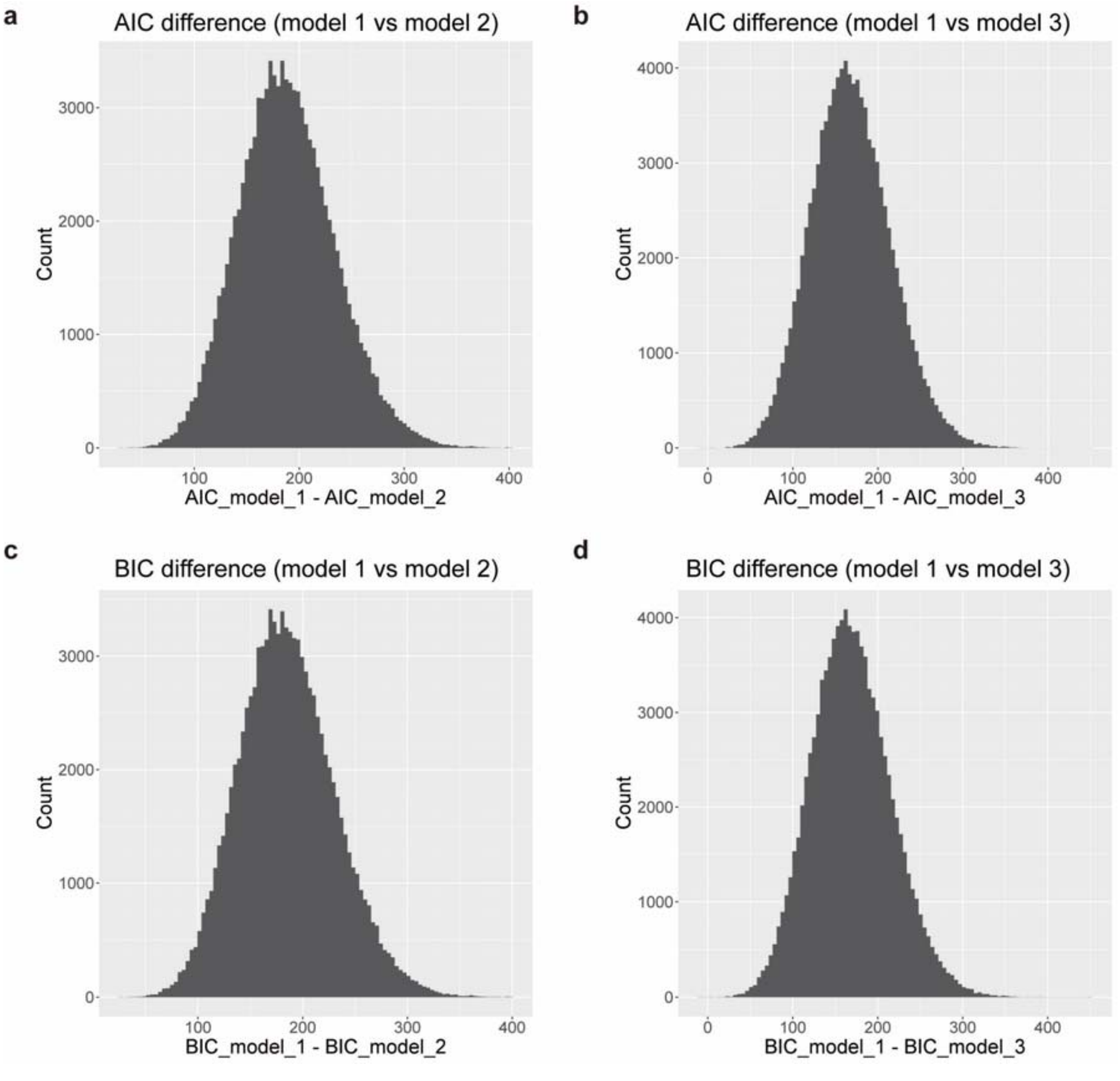


Histogram of bootstrap statistics comparing the Cox proportional hazards model.

**a**, AIC of Model 1 – AIC of Model 2. **b**, AIC of model 1 – AIC of Model 3. **c**, BIC of model 1 – BIC of Model 2. **d**, BIC of model 1 – BIC of Model 3. AIC, Akaike information criterion; BIC, Bayesian information criterion.
